# Parallel and recurrent cascade models as a unifying force for understanding sub-cellular computation

**DOI:** 10.1101/2021.03.25.437091

**Authors:** Emerson F. Harkin, Peter R. Shen, Anish Goel, Blake A. Richards, Richard Naud

**Affiliations:** uOttawa Brain and Mind Institute, Centre for Neural Dynamics, Department of Cellular and Molecular Medicine, University of Ottawa, Ottawa, ON, Canada; Department of Systems Design Engineering, University of Waterloo, Waterloo, ON, Canada; Lisgar Collegiate Institute, Ottawa, ON, Canada; Mila, Montréal, QC, Canada; Montreal Neurological Institute, Montréal, QC, Canada; Department of Neurology and Neurosurgery, McGill University, Montréal, QC, Canada; School of Computer Science, McGill University, Montréal, QC, Canada; Department of Physics, University of Ottawa, Ottawa, ON, Canada

**Author notes:** These authors contributed equally (Emerson F. Harkin, Peter R. Shen). Co-corresponding authors, order decided by coin toss. Email addresses (Blake A. Richards), (Richard Naud).

**Keywords:** Cascade models, Single-cell computation, Dendritic non-linearities, Artificial neural networks

## Abstract

Neurons are very complicated computational devices, incorporating numerous non-linear processes, particularly in their dendrites. Biophysical models capture these processes directly by explicitly modelling physiological variables, such as ion channels, current flow, membrane capacitance, etc. However, another option for capturing the complexities of real neural computation is to use cascade models, which treat individual neurons as a cascade of linear and non-linear operations, akin to a multi-layer artificial neural network. Recent research has shown that cascade models can capture single-cell computation well, but there are still a number of sub-cellular, regenerative dendritic phenomena that they cannot capture, such as the interaction between sodium, calcium, and NMDA spikes in different compartments. Here, we propose that it is possible to capture these additional phenomena using <u>parallel, recurrent</u> cascade models, wherein an individual neuron is modelled as a cascade of parallel linear and non-linear operations that can be connected recurrently, akin to a multi-layer, recurrent, artificial neural network. Given their tractable mathematical structure, we show that neuron models expressed in terms of parallel recurrent cascades can themselves be integrated into multi-layered artificial neural networks and trained to perform complex tasks. We go on to discuss potential implications and uses of these models for artificial intelligence. Overall, we argue that parallel, recurrent cascade models provide an important, unifying tool for capturing single-cell computation and exploring the algorithmic implications of physiological phenomena.

## Introduction

One of the greatest success stories in modern neuroscience is the development of biophysical models of single-cell computation. Though there are still many mysteries to be explored, and new discoveries are still being made, our core understanding of the biophysics of excitable membranes as described by circuit equivalence models, cable theory, and Hodgkin & Huxley-style models has stood the test of time and can reasonably be considered as an accepted theory in neuroscience (Brunel et al., 2014, Herz et al., 2006, Hodgkin and Huxley, 1952, Koch, 2004, McKenna et al., 2014, Rall, 1960). This has provided the foundation for countless computational studies on single-cell computation using detailed models that explicitly incorporate physiological variables including membrane capacitance, ion channels, reversal potentials, etc. Such models have proven very useful for understanding a variety of phenomena, particularly dendritic computation (Poirazi and Papoutsi, 2020, Mainen and Sejnowski, 1996, Schaefer et al., 2003, Vetter et al., 2001, Shai et al., 2015, Psarrou et al., 2014, Gidon et al., 2020, Krichmar et al., 2002, Cook and Johnston, 1997, Gasparini et al., 2004, Ariav et al., 2003, Eyal et al., 2014, Deitcher et al., 2017, Papoutsi et al., 2017). Without these models we would understand much less than we do about how dendrites contribute to computation in neural circuits.

However, due to their complexity, biophysical models are very difficult to link to algorithmic models of neural computation. To some extent, this is part of the dilemma we always face in science, i.e., “How detailed should our models be in order to achieve our scientific goals?” (Poirazi and Pa-poutsi, 2020, Richards et al., 2019). But, one thing that we can say is that it would be beneficial for understanding the functional importance of dendritic computation if we possessed models of intermediate complexity that could capture many of the phenomena of single-neuron computation while still being sufficiently mathematically tractable to use for explaining complex animal behaviour. Moreover, if we could develop such intermediate models we would be better placed to use insights on dendritic computation to inform the development of novel machine learning approaches (Richards et al., 2019, Sinz et al., 2019).

To this end, previous work has explored the use of “cascade models” to capture dendritic computation (Poirazi et al., 2003, Tzilivaki et al., 2019, Kalmbach et al., 2017, Ujfalussy et al., 2018) (but see (Francioni and Harnett, 2021). Cascade models use a cascade of linear and non-linear operations, which are effectively tree-structured, feedforward, multi-layer artificial neural networks (ANNs) (Poirazi and Papoutsi, 2020). Research has shown that these models can capture more variance in single-cell activity than standard point neuron models (which consist of a single linear operation and non-linear activation function) (Poirazi et al., 2003, Tzilivaki et al., 2019, Ujfalussy et al., 2018). Thus, cascade models treat <u>individual</u> neurons as deep feedforward ANNs in order to capture single-cell computation with a mathematically tractable model (Figure 1A). However, such models still have not been compared to many facets of dendritic computation, including calcium spikes and N-methyl-D-aspartate (NMDA) receptor mediated plateaus.

**Figure 1:**
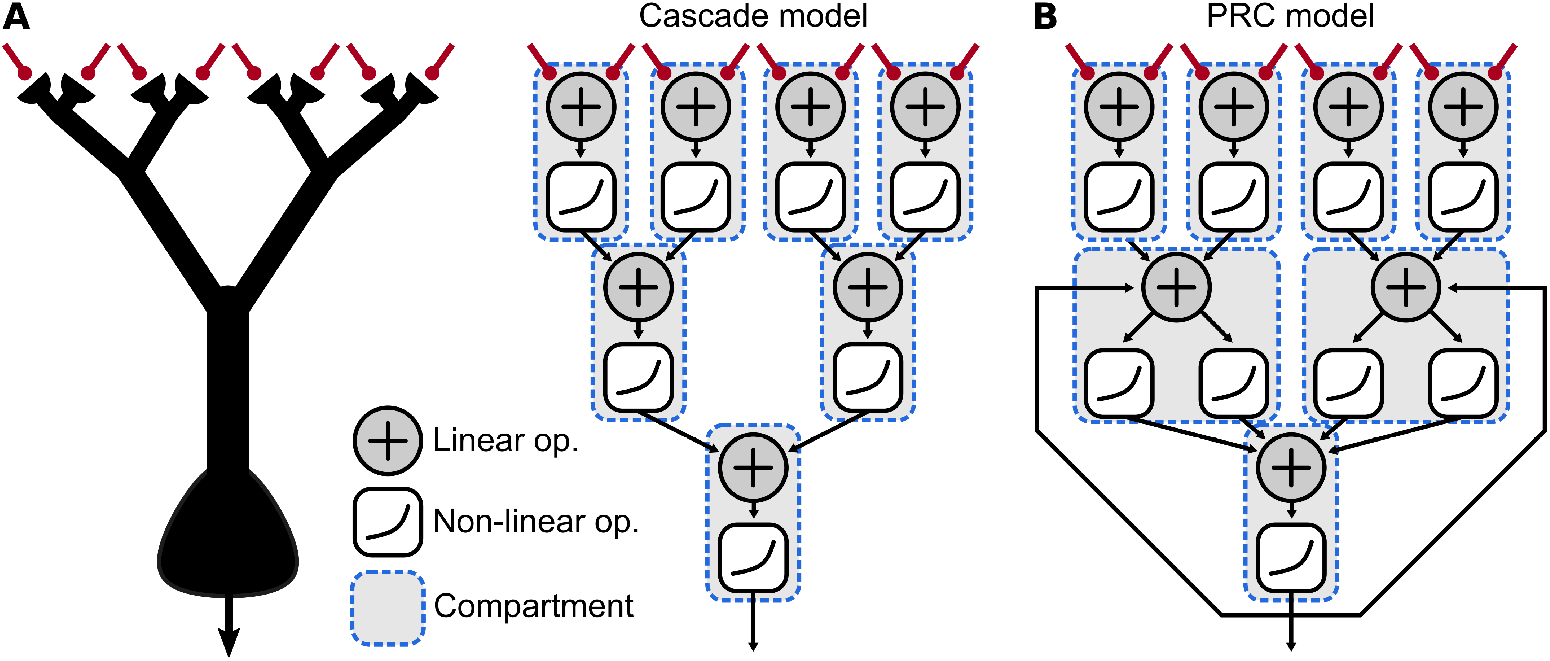
Illustration of cascade models and PRC models: **A** Dendritic computation can be modelled using a cascade of linear and non-linear operations (Poirazi et al., 2003, Tzilivaki et al., 2019, Ujfalussy et al., 2018). **B** Cascade models can be extended with the use of parallel pathways and recurrence in the operations, which can enable the modeling of more complicated regenerative phenomena.

Here, we show that it is possible to capture these phenomena using **parallel and recurrent cascade models** (PRC models), i.e. models that use a set of parallel cascades of linear and non-linear operations that are recurrently connected to one another. This is equivalent to treating individual neurons as multi-layer, recurrent ANNs (Figure 1B). We show that these PRC models can successfully reproduce a number of experimentally observed regenerative phenomena in dendrites, all while being mathematically tractable. Moreover, because these models are fully differentiable, we show that they can easily be incorporated into machine learning approaches that utilize gradient descent for model optimization (Richards et al., 2019). This opens the door to exploring the possibility that dendrites and dendritic computation may provide important inductive biases for brains that could be mimicked by artificial intelligence (AI) (Richards et al., 2019, Sinz et al., 2019). Thus, we conclude this paper by providing some speculation as to the possible advantages for AI of dendritic PRC models. Altogether, this work helps to lay the ground for better integration between our well-developed understanding of the biophysics of neural computation and our ever increasing understanding of algorithms for complex behaviour.

## The use of recurrent cascade models to capture single-cell computation

Models made of a cascade of linear-nonlinear (LNL) operations have had a long history in systems neuroscience, where such models were conceptually implied by early work on retinal ganglion cells (Kuffler, 1953) and cortical cells (Hubel and Wiesel, 1968). These models consist of a linear filter of the stimulus followed by a nonlinear readout to generate predictions of a nonnegative observable. Important refinements to improve the accuracy of these models were the inclusion of spike-triggered adaptation (Pillow et al., 2005, Truccolo et al., 2005), the composition of linear-nonlinear (LNL) operations in a cascade (Vintch et al., 2015, McFarland et al., 2013), and the addition of recurrent interactions between linear-nonlinear subunits (Pillow et al., 2008, Truccolo et al., 2011). Together, these various extensions are described by an LNL subunit with multiple possible types of interconnection motifs. The LNL subunit is composed of a linear-nonlinear operation with feedback from the output of the nonlinearity to capture spike-triggered adaptation (Fig. 2A; Methods 1.1). Wiring between subunits can create multiple types of motifs including a strictly feedforward cascade, a common input to two units having possibly different filters or nonlinearities (parallel feature processing, Fig. 2B), or recurrent interconnections (Fig. 2C). In most systems neuroscience applications, the output of the nonlinearity is the Poisson intensity, used to capture the stochastic spike-timing responses of real neurons. Our approach with PRC models here is slightly different, as we will consider the output of the nonlinearity to represent deterministic voltage excursions. Also, in most systems neuroscience applications, the filters and nonlinearities may arise from a large number of possible – yet undefined – mechanisms. These are typically thought of as interactions within and between cells, but may also include dendritic computations (Taylor et al., 2000). In order to better understand the cellular mechanisms underlying such information processing, we focus on the application of a deterministic LNL framework within a single cell.

**Figure 2:**
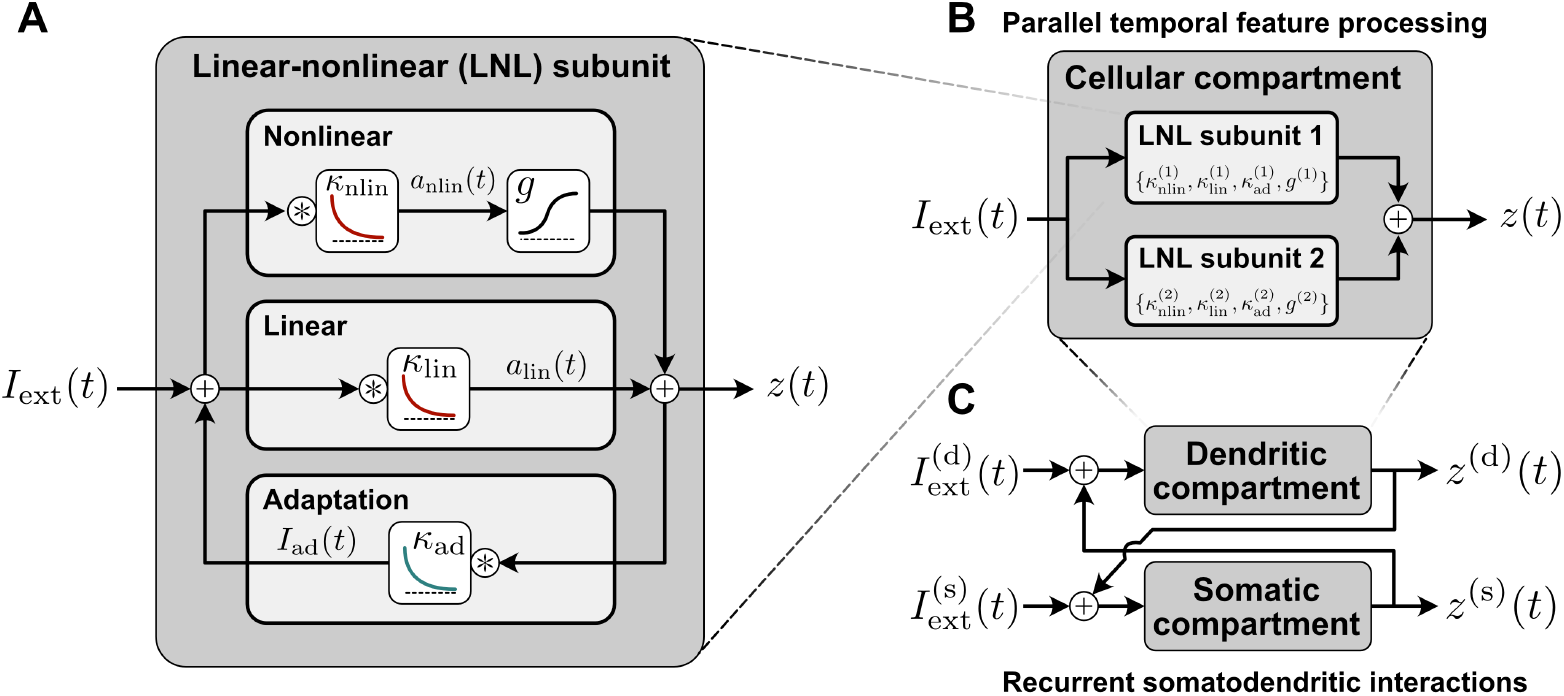
Linear-nonlinear operations and interconnection motifs. **A** The linear-nonlinear (LNL) subunit. This architecture combines linear filtering of an input *I*_ext_(*t*) (middle) with a nonlinear readout (top, *g*) and feedback (bottom, *κ*_ad_), which together flexibly capture the passive filtering effects of neuronal membranes, the nonlinear effects of voltage-dependent ionic conductances, and adaptation. The contributions of each of these effects to the output *z*(*t*), which may loosely represent neuronal voltage or a spiketrain, can be tuned via the parameters of the filters *κ*_nlin_, *κ*_lin_, and *κ*_ad_, and the choice of nonlinear function *g*(·). **B** Multiple LNL subunits arranged in parallel can model the effects of multiple ionic conductances in a single cellular compartment. **C** Recurrent connections between model compartments, each composed of one or more LNL subunits, capture interactions between cellular compartments.

### Somatic spikes

Since the pioneering work of Richard Stein (Stein, 1965), the leaky integrate- and-fire (LIF) model has become the workhorse of interrogations of information processing of large numbers of interconnected neurons. In itself, the LIF can be seen as a special case of the LNL subunit. When a depolarizing current is injected into an LIF model, it first passes through the membrane filter and produces a voltage (leaky integration; a linear operation) which is eventually reset to a lower value if it reaches a threshold (firing; a nonlinear operation). This leaky integrate-and-fire behaviour can be captured by an LNL subunit with a filter that corresponds to the membrane filter of an LIF model, together with a Heaviside nonlinearity that is triggered exactly at spike threshold and a Dirac delta-shaped adaptation filter which resets the voltage to a lower value. Such deterministic LNL models can capture both the time-dependent membrane voltage response and the spike timing responses to complex inputs (Jolivet et al., 2006). Adding multiple timescales to the kernel for spike-triggered adaptation makes these models highly predictive of the responses of a variety of cell types (Jolivet et al., 2008, Mensi et al., 2012, Pozzorini et al., 2013, Kobayashi et al., 2009, Teeter et al., 2018). Furthermore, considering smooth nonlinearities and surrogate gradients can ensure that the LNL unit remains differentiable (Neftci et al., 2019).

In addition it is worth noting that since LNL subunits are a special case of PRC models, and LIF models are a special case of LNL models, LIF models are, in fact, a restricted class of PRC models. As such, they can capture some of the phenomena that the broader class of PRC models are capable of capturing (Weber and Pillow, 2017, Gerstner et al., 2014). Part of our contribution here is to illustrate some of the more complicated sub-cellular phenomena that cannot be recapitulated with a pure LIF model, such as dendritic non-linearities and coincidence detection (see sections 2.3-5, below).

### LNL models for dendritic spikes

To circumscribe a systems-level function for dendritic computation, many studies have focused on the role of intrinsic nonlinear dendritic operations— first using models of dendritic trees in a stationary state (Mel, 1993, Poirazi and Mel, 2001, Poirazi et al., 2003), then using models capturing the dynamics of dendrites and their intrinsic nonlinear properties (Legenstein and Maass, 2011, Kalmbach et al., 2017, Ujfalussy et al., 2018). These contributions are examples of what we call PRC models because they involve a cascade of nonlinearities, but they leave out the recurrent motifs. Also, most have not considered the parallel processing introduced by Ujfalussy et al. (2018) (Ujfalussy et al., 2018). Recurrence was, however, part of other efforts focusing on simplified models of the interaction between somata and dendrites (Pinsky and Rinzel, 1994, Urbanczik and Senn, 2014, Naud et al., 2014, 2017). Thus our goal in this section is to unify these complementary perspectives by presenting data showing that PRC models can reproduce <u>qualitative</u> features of sub-cellular physiological phenomena, such as dendritic spikes, coincidence detection, etc.

In comparison to the action potentials generated in the proximity of the cell body, the spikes observed in dendrites display less stereotypical amplitudes and durations (Golding and Spruston, 1998, Larkum and Zhu, 2002, Smith et al., 2013, Schiller et al., 2000). These features pose a problem for the LIF framework, but they are captured naturally by the LNL framework. In Figure 3, we illustrated the response of a LNL subunit to noisy inputs and to short pulses. For a fast filter preceding a sharp nonlinearity, the model produced short spikes on top of a noisy voltage trace. The short spikes arise when the low-pass filtered input reaches the activation threshold of the nonlinear readout. Sometimes, the fluctuation is only able to activate a fraction of the nonlinearity, which produces spikes of variable amplitudes. Increasing the mean of the input makes those spikes more frequent as there are more chances that the random fluctuations reach the activation threshold of the nonlinearity. Thus in the configuration where a sharp sigmoidal nonlinearity is fed by a linear filter that is considerably faster than the membrane filter, we observe variable amplitude spikes akin to dendritic sodium spikes. We note that a very similar model architecture was able to be reproduce with great precision the response to noisy dendritic inputs in the presence of dendritic sodium spikes (Kalmbach et al., 2017).

**Figure 3:**
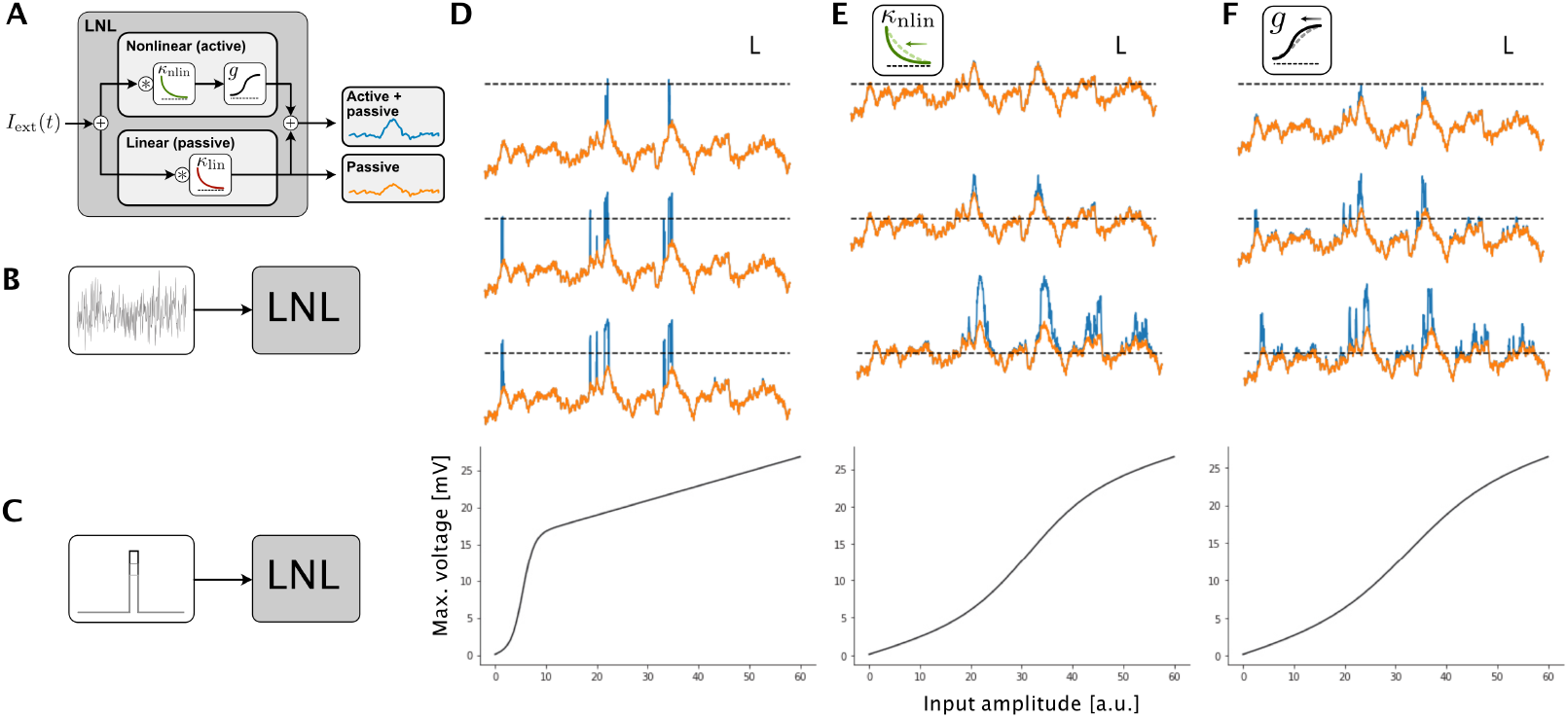
Effects of changing parameters in the linear-nonlinear model. **A** Schema of the LNL model. An external input undergoes two parallel processing steps: a linear (passive) filter and a linear filter (possibly distinct from passive) followed by a nonlinear readout. The nonlinear processing step (top) produces a nonlinear component (blue) which is added to a passive component (orange). As input to the model, we consider both **B** a noisy time-dependent signal representing a bombardment of asynchronous synaptic currents, and **C** a short pulse. **D, top 3 traces** Response to noisy inputs having three different baselines (lower baseline is topmost trace,dashed line represents the activation threshold of the nonlinearity). The nonlinear (blue) is added to the passive component (orange). **D, bottom** Maximum amplitude of response as a function of pulse amplitude (bottom). **E** Same as D but using a model with an increased timescale of the linear filter. **F** Same as D but with a model having a decreased sensitivity (steepness) of the nonlinearity. Scale bars correspond to 10 ms and 5 mV.

Next, we examined the effects of changing the parameters in the LNL subunit (Fig. 3E). We began by increasing the timescale of the filter preceding the nonlinearity. This reduced the number of suprathreshold fluctuations, and when a fluctuation in the low-pass filtered input did cross the activation threshold it tended to stay for a longer time period. This produced less frequent but longer spikes, akin to calcium spikes (Larkum et al., 1999, Larkum and Zhu, 2002, Magee and Carruth, 1999, Xu et al., 2012) or, with an even longer timescale, NMDA spikes (Schiller et al., 2000). As a consequence of changing the filter, the aspect of the nonlinearity that can be observed when presenting a pulse input is altered, and appears more graded. When, instead of changing the timescale of the linear filter, we only changed the gain of the nonlinearity (Fig. 3F), then the spikes had a more variable amplitude and duration. Changing the timescale of the filter and the steepness of the nonlinearity had a similar effect on the maximum voltage observed in response to a pulse input. This can be explained by the fact that in response to a short pulse, the voltage increases linearly with a slope given by the timescale of the filter (Gerstner et al., 2014) and thus acts as a multiplicative factor identical to the steepness of the nonlinearity. In these simulations, we have not included an adaptation-like recurrence, although the formalism can include this mechanism. Thus, altogether, by changing the parameters of the LNL model, we can simulate some basic electrophysiological features of various types of dendritic spikes.

### Dendritic sodium spikes

To test whether a PRC model can capture other features observed in electrophysiological recordings, we focused on experimental findings reported by Golding and Spruston (1998) (Golding and Spruston, 1998) pertaining to dendritic sodium spikes. In one of the experiments reported (Fig. 4), recordings were made simultaneously in a proximal dendrite and in the soma. A variable-amplitude synaptic-like stimulus was injected in the dendrite. The recordings showed that in one dendrite, low input amplitudes initiated a spike in the soma which produced a back-propagating action potential in the dendrite and at high amplitudes initiated a spike in the dendrite <u>before</u> triggering an action potential in the soma. In another dendritic recording, a low amplitude stimulus initiated a dendritic spike unaccompanied by an action potential at the soma, and only a large input produced an action potential at the soma. We found that we can reproduce these phenomena by changing the parameters of two LNL subunits wired in a recurrent fashion (Fig. 4A and C). To capture how different recordings initiated spikes preferentially in the dendrite or the soma, we varied the relative threshold of activation of the dendritic and somatic nonlinearities.

**Figure 4:**
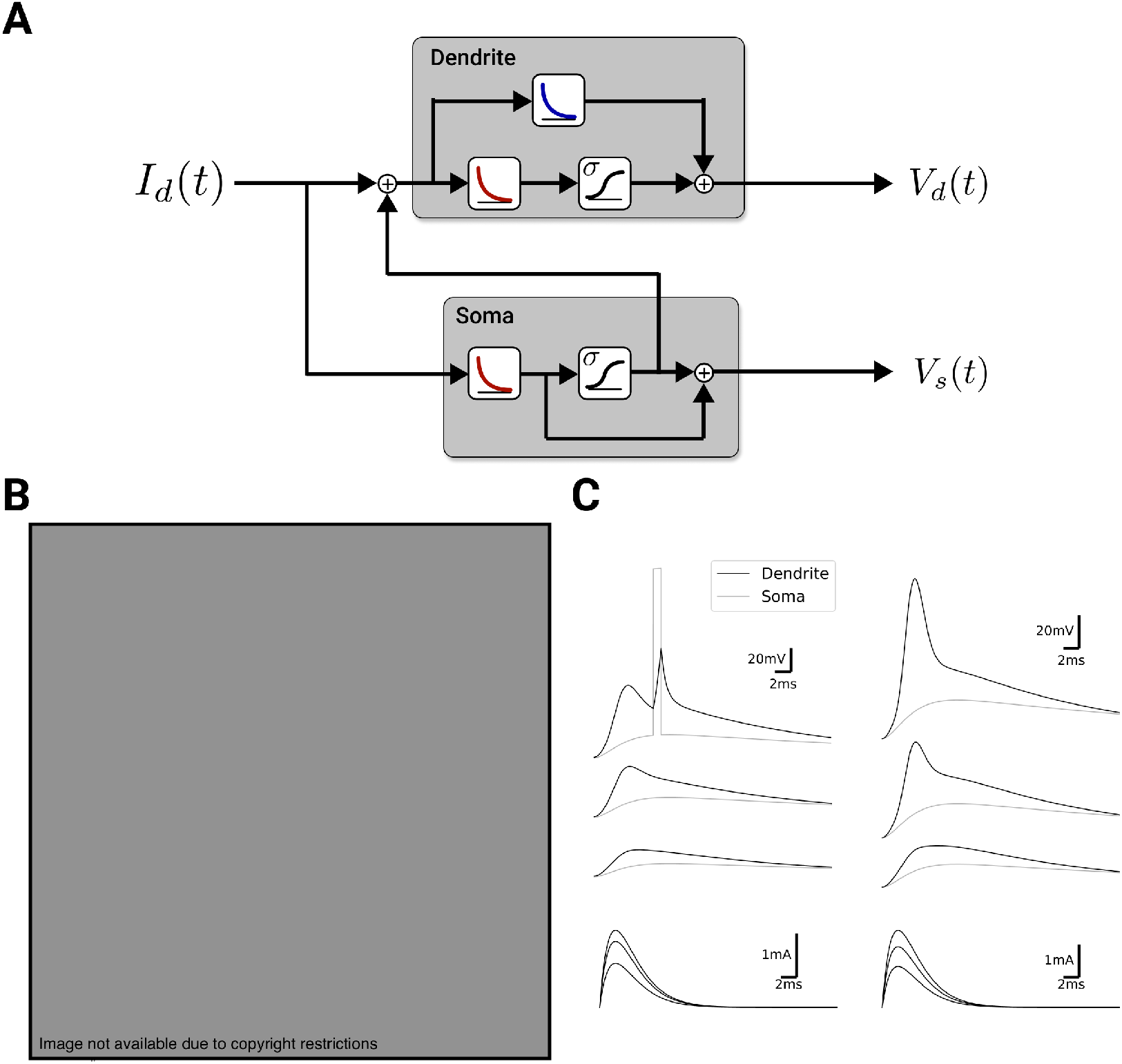
A recurrent motif of linear-nonlinear models for the dendritic sodium spikes. **A** Schematic of the model: A dendritic current (*I_d_*) impinges on two LNL subunits with recurrent interactions, one corresponding to a dendrite, another corresponding to the soma. When the somatic compartment reaches the threshold of its nonlinearity, a spike in the form of a square pulse is added to the dendritic current. **B** Experimental data showing injection of synaptic-like pulses of varying amplitudes in the dendrite (topmost traces have lowest input), where the voltage in the dendrite (thick trace) and soma (thin trace) are shown. Two exemplars are shown in different columns. Reproduced from Golding et al. (1998) Fig. 1 (Golding and Spruston, 1998). **C** Model responses showing three amplitude levels of a synaptic-like input in the model shown in A. To reproduce the exemplars in B the model in the right column has a lower threshold for the dendritic nonlinearity and a higher threshold for the somatic nonlinearity.

### Dendritic NMDA spikes

Next we considered the electrophysiological recordings of NMDA spikes reported in Schiller et al. (2000) (Schiller et al., 2000) (Fig. 5). We focused on the threshold input-pulse amplitude to trigger an NMDA spike which was lowered by the addition of blockers of sodium and calcium ion channels (TTX and cadmium). This observation suggested that the nonlinear effects of sodium and calcium ion channels participated in the initiation of the NMDA spikes. Since calcium and sodium ion channels are characterized by distinct timescales, we considered a parallel connectivity motif shown in Fig. 5A. We chose the filter of the sodium LNL to be fast (5 ms), the filter of the calcium filter to be slower (40 ms), and the filter of the NMDA LNL to be even slower (80 ms). Simulating the response of this model to pulse inputs produced a long depolarization that was clearly initiated by contributions from sodium and calcium (Fig. 5B). To simulate the blockade of these mechanisms by TTX and cadmium, we reduced the amplitudes of their corresponding nonlinearities to zero, which prevented the occurrence of spikes for a range of input amplitudes (Fig. 5C). The effect on the initiation threshold of removing the nonlinearity in the PRC model mimicked that of pharmacological manpulations (Fig. 5D-E).

**Figure 5:**
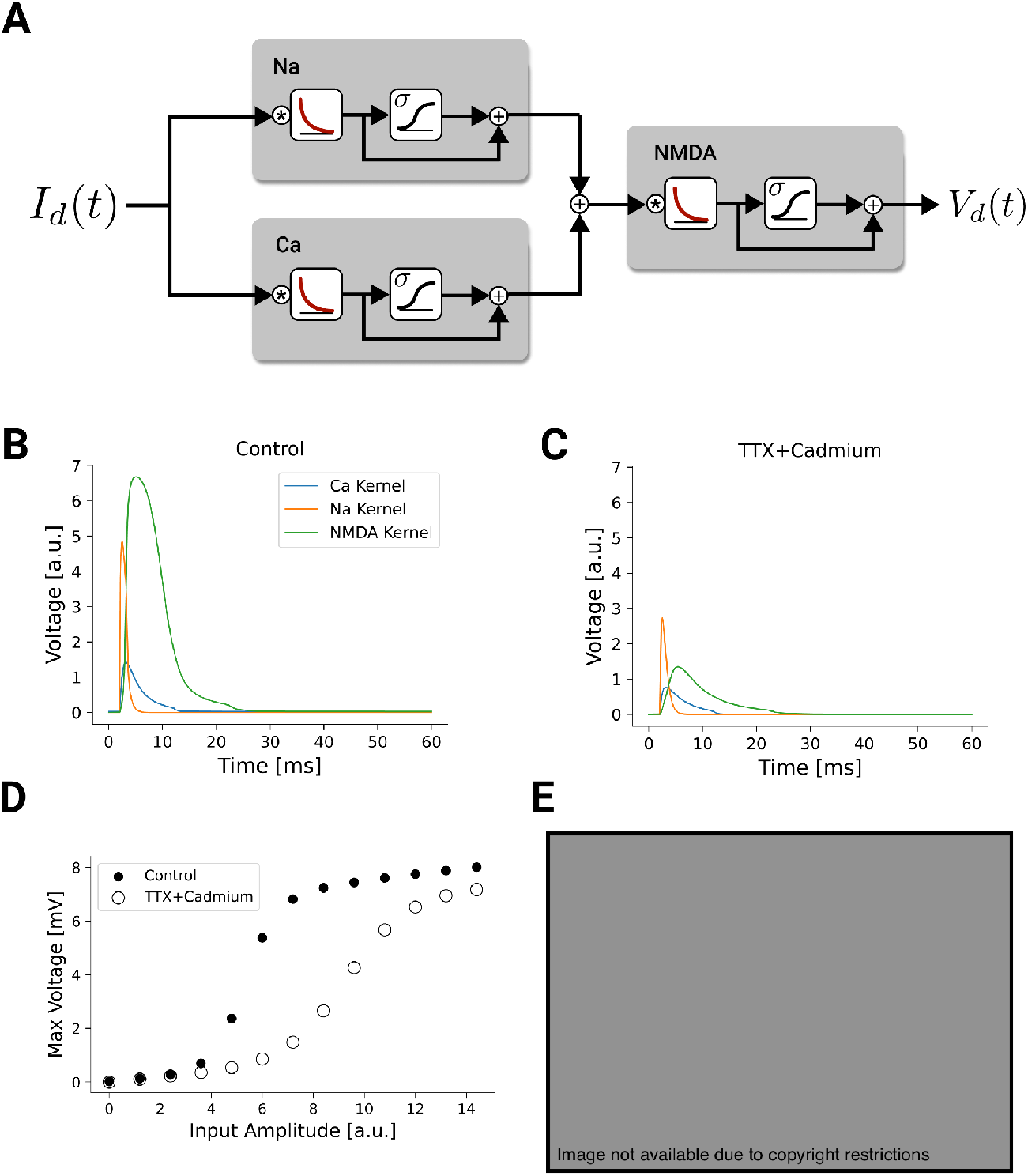
Linear nonlinear model of a NMDA spike as a combination of cascade and parallel processing. **A** Schematic of the model: current impinging on the dendrite is passed through two LNL operations in parallel before feeding into another LNL operation to produce an NMDA spike. **B** Response of the model to a supra-threshold pulse input showing the calcium (blue) sodium (orange) and NMDA (green). **C** The response to the model with the same pulse input as in B but with the nonlinear component of sodium and calcium set to zero, simulating the application of TTX and Cadmium. **D** Maximum voltage as a function of the amplitude of the input pulse for the model in B (full circles) and the model in C (open circles). **E** Experimental recordings of peak membrane potential as a function of stimulation power in control (full circles) and the presence of calcium and sodium ion channel blockers (TTX and cadmium, open circles). Figure reproduced from Schiller et al. (2000) Fig. 3c (Schiller et al., 2000).

### Dendritic calcium spikes

We then considered how bidirectional interactions between somatic spiking and calcium spikes can be captured by a PRC model. The tuft dendrites of cortical pyramidal cells express a high density of voltage-gated calcium channels which produce dendritic plateau potentials when sufficiently strong inputs are injected into the soma and tuft dendrites simultaneously (Larkum et al., 1999, Magee and Carruth, 1999). These dendritic plateau potentials mediate burst firing at the soma, producing coincidence detection and modulating the gain of somatic responses to peri-somatic input. To capture these effects in our PRC framework, we created a model with two recurrently-connected compartments, corresponding to the soma and apical tuft dendrites (Fig. 6A). Appropriately tuned filtering and nonlinear operations in each compartment (see Methods 1.4) caused the somatic compartment to emit single spikes when input was delivered to the soma alone and intermittent bursts when input was delivered to both compartments simultaneously (Fig. 6B). Inputs delivered to either compartment alone evoked small responses in the dendritic compartment, while simultaneous inputs to both compartments evoked burst-associated plateau potentials in the simulated dendrite. In cortical pyramidal neurons, dendritic inputs produce somatic bursts most potently when they are delivered just before or at the same time as somatic input, creating an asymmetric coincidence-detection effect (Larkum et al., 1999). Injecting a synaptic-like pulse into the dendritic compartment of our model evoked a dendritic plateau potential and somatic burst only when it preceded or arrived at the same time as a somatic input pulse (Fig. 6C), recapitulating the coincidence-detection effect observed in pyramidal neurons (Larkum et al., 1999). Dendritic input to our model also modulates somatic gain by increasing the number of spikes evoked by a given input (Fig. 6D), consistent with an effect of dendritic input on gain observed experimentally (Larkum et al., 2004). Together, these simulations add to previous efforts (Naud et al., 2014) in showing that the PRC framework is able to capture features of dendritic excitability and somatodendritic interactions.

**Figure 6:**
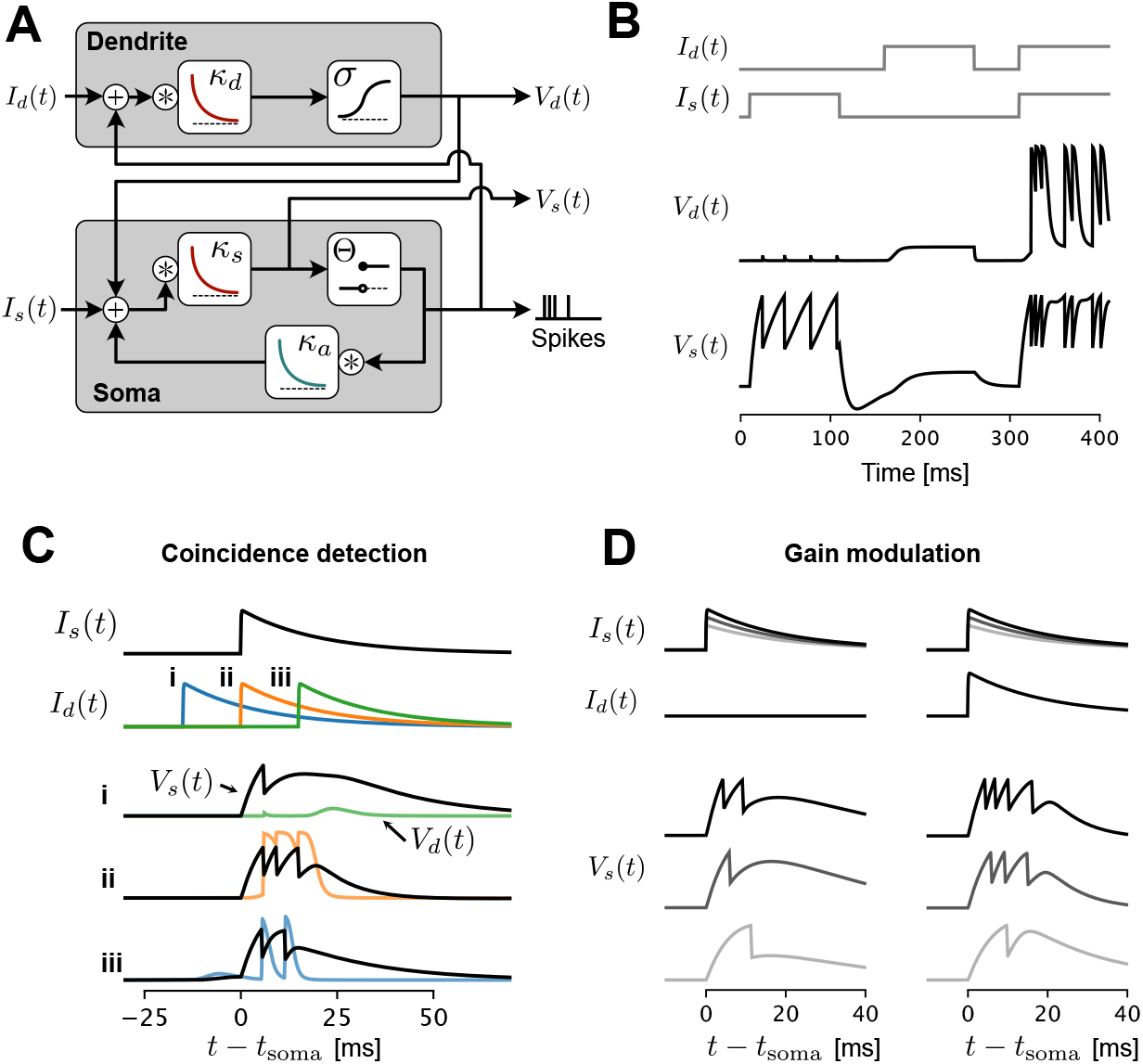
A recurrent cascade model of the interaction between the back-propagating action potential and calcium spikes. **A** Schematic of the model: external current impinging on the dendrite (*I_d_*) and soma (*I_s_*) each engage a LNL operation. The output of the nonlinearities is combined to the input current of the other unit, forming a recurrent interaction. **B** Step current injections in the soma alone, the dendrite alone and then in both compartments simultaneously produces regular bursting only for the conjunction of inputs. **C** A strong synaptic-like current pulse is injected in the both the soma (*I_s_*) and the dendrite (*I_d_*). Three simulations are shown for three relative timings of the dendritic input (i blue, ii orange, iii green). Responses for the somatic (*V_s_*, black traces) and dendritic (*V_d_*, color traces) compartments are overlaid for each condition. Somatic spikes are denoted by a clear reset but the full depolarization is not shown. A burst of action potentials arise from the near-coincident condition (ii, orange) recapitulating experimental observations in pyramidals of the cortex (Larkum et al., 1999). **D** Responses to increasing amplitudes of synaptic-like input to soma in the absence of dendritic input (left) and in the presence of a concomitant input in the dendrite (right), a simulation of the gain modulation property of dendritic input reported experimentally (Larkum et al., 2004).

### Potential applications of recurrent cascade models for learning theory

Aside from the additional capabilities to capture biological phenomena in dendrites that we have illustrated here, PRC models may have relevance for machine learning applications. Notably, thanks to the use of differentiable computational graphs (see (Zenke and Ganguli, 2018) for an approach to making our somatic units differentiable), a PRC model can be incorporated into any artificial neural network model and trained with gradient descent (Ujfalussy et al., 2018, Jones and Kording, 2021). As such, PRC models open the door to investigating whether sub-cellular dendritic computations have any potential utility for improving machine learning applications. The answer to this question will depend, in large part, on whether dendritic computations can provide important inductive biases for an artificial neural network (Bird and Cuntz, 2020, Jones and Kording, 2021).

### Networks of PRC models can be trained using gradient descent

To illustrate that complex PRC models can be incorporated into multilayer artificial neural networks and while remaining trainable via gradient descent, we next trained an artificial neural network containing a single hidden layer made of four PRC neurons to solve a simple binary classification problem: memorizing labels associated with random patterns of synaptic input (Fig. 7A). Before training, our network correctly predicted the label (“green” or “purple”) that had been randomly assigned to each randomly-generated pattern of synaptic input ~50 % of the time, no better than chance. Adjusting the synaptic weights in the direction of decreasing error using back-propagation through time (with a surrogate gradient for the somatic spiking function; see Methods) gradually improved the prediction accuracy to well above the chance level (>75%, Fig. 7C, D), indicating that the network had successfully learned a mapping from synaptic input patterns to class labels. We repeated the training procedure with different types of hidden units (Fig. 7E), ranging from an integrate-and-fire unit without a dendrite (model ①) to the full PRC model (model ⑤), and including recurrently-connected compartments without paralel processing (as in Fig. 6). For the models with a dendrite (②–⑤ in Fig. 7), two independent sets of weights were used, such that each input unit connects with one trainable weight to the soma and with another trainable weight to the dendrite. Also, for the more complex models (②–⑤ in Fig. 7), we fixed as hyper-parameters the connection strength between the somatic and dendritic compartment as well as the time scale of the filters (see Methods). We found that, with the exception of the models with the parallel motif, all models trained well and achieved above 80% accuracy within 2000 epochs, demonstrating these models are trainable using standard techniques.

**Figure 7:**
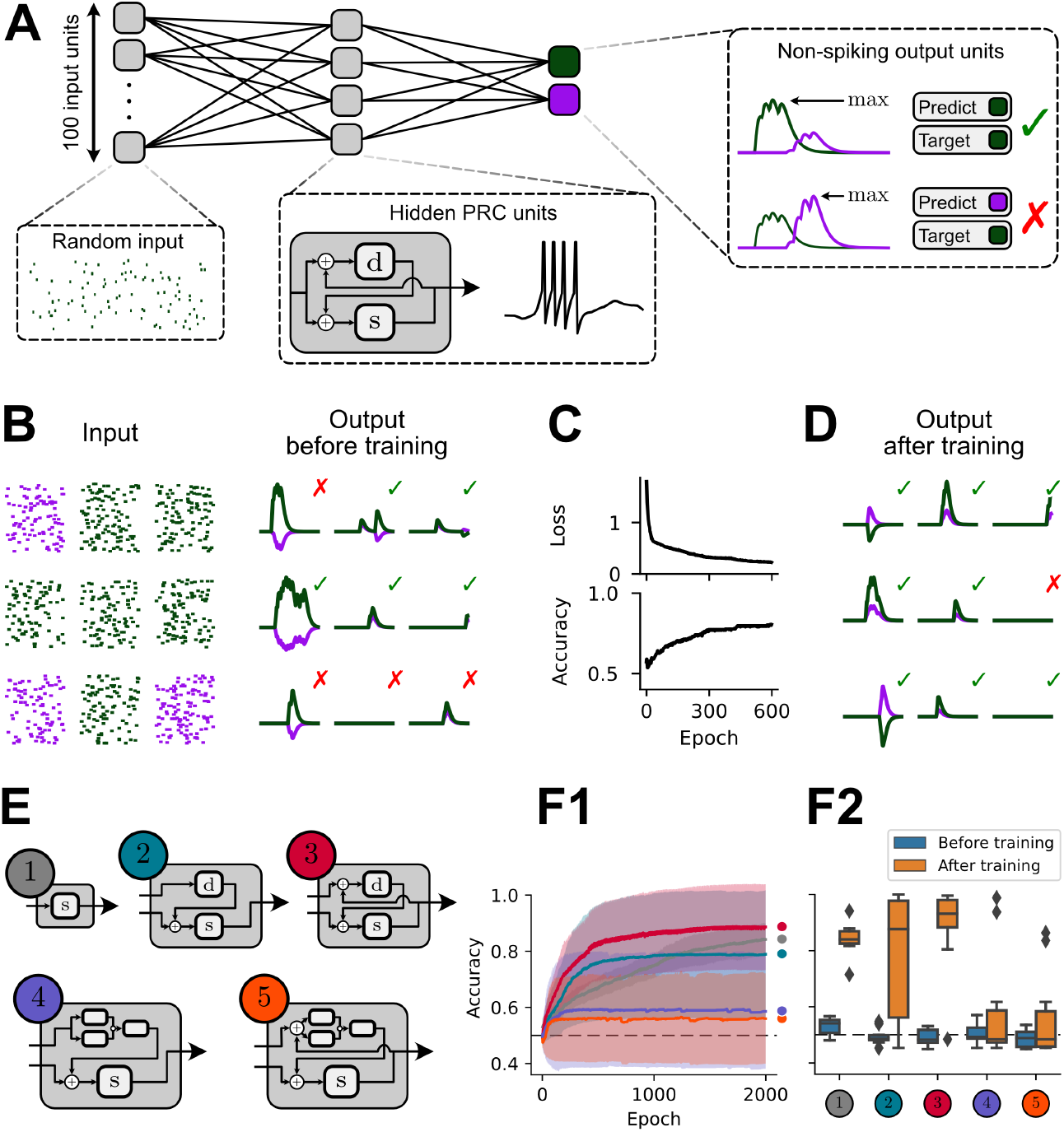
PRC neurons in a network can be trained to memorize patterns of synaptic input. **A** Network architecture with PRC units in the hidden layer. 100 input units send synaptic input to a hidden layer composed of four PRC neurons with recurrently-connected somatic and dendritic subunits (as shown in inset and Fig. 6A), which in turn send synaptic input to an output layer consisting of two non-spiking single-compartment neurons. Synaptic input patterns are classified as “green” or “purple” based on the maximum voltage attained in the output layer. **B** Randomly-generated patterns of synaptic inputs (left) are incorrectly classified approximately 50 % of the time by a PRC network with randomly-initialized synaptic weights. **C, D** Training synaptic weights using back-propagation through time allows the network to memorize labels associated with specific input patterns, resulting in improved predictions. The mean negative log likelihood loss is shown (top). A surrogate gradient was used for the Heaviside nonlinearity in the somatic compartment of hidden PRC neurons (see Methods) in order to train the input weights. **E** Different architectures of the PRC unit used in the hidden layer: ① one-compartment neuron, ② two-compartment neuron with no recurrent connection from the soma to dendrite, ③ two-compartment neuron with recurrent connections between soma and dendrite (as in 2C and 6), ④ two-compartment neuron with parallel processing in the dendritic compartment (as in 5), and (⑤ two-compartment neuron with both parallel processing and recurrent somatodendritic connections. **F1** Mean ± SD accuracy during training for *N* = 10 randomly-initialized networks of each type. Dashed lines indicate chance-level accuracy. **F2** Training set accuracy before and after training is shown for the different models.

Importantly, as with any machine learning model, increasing PRC model complexity increases the number of parameters to be trained and hyperparameters to be tuned. Good machine learning practice involves a systematic methodology for tuning the hyper-parameters (Bergstra et al., 2011, Feurer and Hutter, 2019). Here, however, the particular choice of hyper-parameters was based on our comparison with electrophysiological observations. With these hyper-parameters, good accuracy using the parallel motif was achieved but only in a small portion of the initializations. It is possible that the slow time constant used in the parallel subunits tended to prevent these models from achieving good accuracy. Thus, we have demonstrated that various forms of PRC models can be trained on a memorization task and that combining these models with techniques for hyper-parameter optimization (Bergstra et al., 2011, Feurer and Hutter, 2019) opens the door to comparing model architectures on the basis of training efficacy.

Because few problems solved by machine learning systems or the brain are likely to closely resemble memorization of random patterns of inputs, we next assessed whether our PRC models could be trained to solve a more realistic problem: classification of spoken digits. The Heidelberg digits dataset contains spiketrains produced by a detailed model of the inner ear in response to audio recordings of the numbers zero to nine spoken by multiple people in two different languages (English and German; Fig. 8A). As a benchmark for assessing the generalization performance of spiking neural network models, the Heidelberg dataset is divided into a standard set of training and test examples, where the test examples include digits spoken by two individuals not included in the training set. To solve this more complex task, we created a set of larger multi-layered PRC network models containing a hidden layer with 200 PRC units in addition to the 700 input units and 20 output units required by the structure of the dataset (Fig. 8B). Training multi-layered networks containing the same set of five PRC models shown in Fig. 7E for 200 epochs consistently improved the training set prediction accuracy from the chance level of 5% to ≥95% and as high as 99.5% for the network containing PRC units which capture the effects of backpropagating action potentials (i.e., ③ in Figs. 7 and 8; analogous to Fig. 6; all values are averages across *N* = 3 randomly-initialized networks; Fig. 7C). The multi-layered PRC networks successfully generalized to new examples, achieving a test set accuracy of 54% to 70%, comparable to that previously reported using networks of LIF neurons (Cramer et al., 2020). The prediction accuracy could be further improved by systematic selection of model hyperparameters, including PRC neuron architecture (Bergstra et al., 2011, Feurer and Hutter, 2019). Importantly, these preliminary results provide a clear proof-of-concept that multi-layered networks containing PRC neurons can be trained to solve complex tasks.

**Figure 8:**
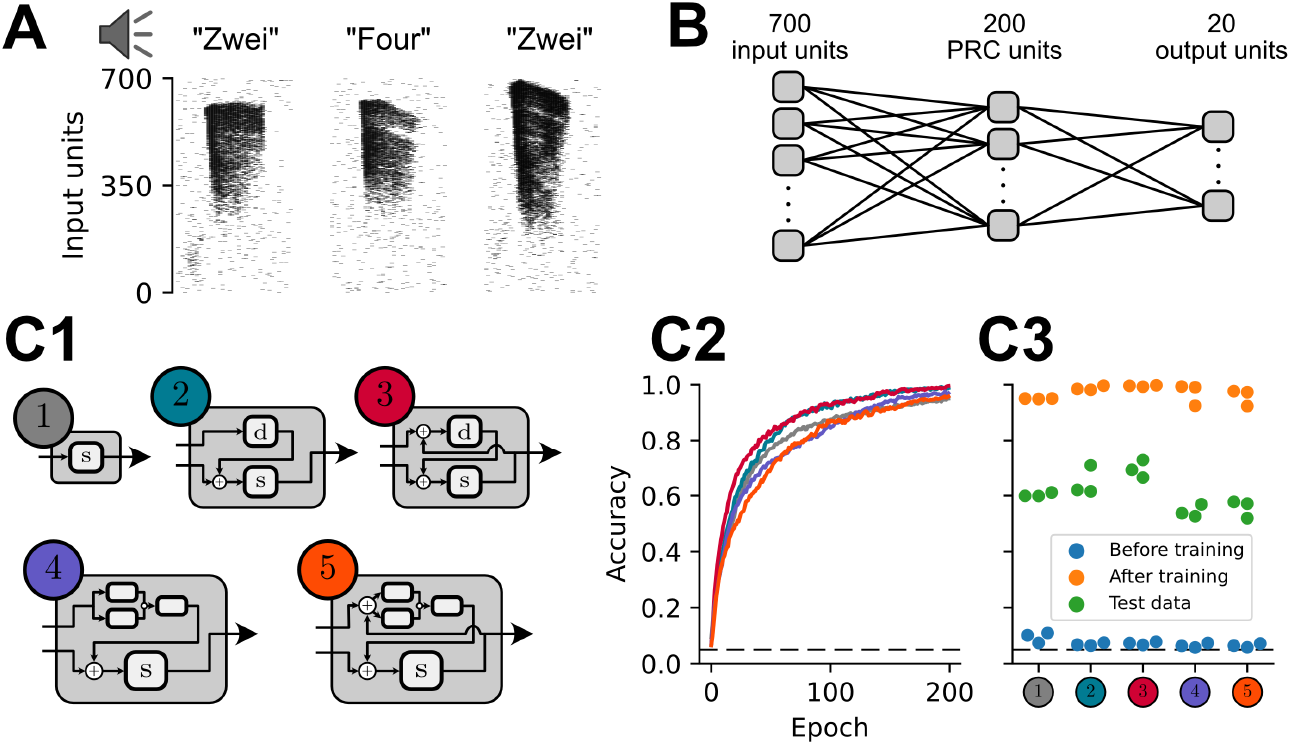
Multi-layered networks of PRC neurons can be trained to recognize spoken digits, and generalize to sound samples from new speakers. **A** Training examples from the Heidelberg spoken digits dataset (Cramer et al., 2020). Audio recordings of digits spoken in English and German were transformed into spiketrains using a detailed model of the inner ear. **B** Multi-layered PRC network architecture, an expanded version of the architecture shown in Fig. 7. **C** Performance of multi-layered PRC networks. Hidden unit architectures (C1) are the same as in Fig. 7E. C2 shows the mean training accuracy across *N* = 3 randomly-initialized networks. Dashed lines indicate chance-level accuracy. C3 shows test set accuracy (green) as well as initial and end training set accuracy for the different models.

### Parallel-recurrent cascade models as architectural inductive biases

One of the realities that any machine learning system faces is that no learning agent can be efficient at learning all types of problems and/or tasks. Due to the No Free Lunch Theorem for Optimization (Wolpert and Macready, 1997), we know that there is always a trade-off: for a learning system to achieve good performance in one set of problems, it must sacrifice its performance in other sets of problems. When an engineer introduces some design components into an optimization system that help to bias a learning system towards a particular set of problems, we refer to these components as “inductive biases”. Inductive biases are key to developing useful machine learning systems, because without appropriate inductive biases, learning systems will provide mediocre performance on all tasks, as opposed to superior performance on the subset of tasks that we may care about (Hessel et al., 2019, Goyal and Bengio, 2020). For example, if an engineer creates a machine learning system that is built with an inductive bias to seek out relationships between discrete objects, then that can help the system learn about spatio-visual object relations such as “there is a blue ball to the left of a red cube” (Battaglia et al., 2018, Santoro et al., 2017).

These insights about the importance of inductive biases actually helped to lay the foundations for the modern deep learning approach in AI (Bengio et al., 2007, Goyal and Bengio, 2020). The early proponents of deep learning proposed that the set of problems we most care about in AI are those that humans and/or animals are good at, e.g. image processing, motor control, language comprehension, etc (Bengio et al., 2007). Given this, they argued that machine learning researchers should seek inspiration from real brains in order to find appropriate inductive biases for AI (LeCun et al., 2015). At the time, the principal inductive bias that these researchers were interested in was network <u>depth</u> (hence the name “deep” learning). They believed that the macroscopic architecture of the brain, with multiple stages of processing involved in sensorimotor transformations, provided inductive biases to promote hierarchical representations, which was proposed to solve the type of problems where humans and animals excel (Bengio et al., 2007). In hindsight, it appears that their intuitions were largely correct: deep ANNs have consistently outperformed other types of machine learning approaches in exactly the sort of problem/task domains that humans and animals excel at (LeCun et al., 2015). Moreover, theoretical analyses have provided some explanations for why ANNs with deep architectures are particularly well suited to such applications (Lee et al., 2017, Li and Sompolinsky, 2020).

There is a broader, two-fold point within the story of deep learning and inductive biases. First, it is clear now that the inductive biases of an ANN are determined in large part by architecture, i.e. how the linear and non-linear operations are arranged in a computational graph within the ANN. This is because architecture determines both how information flows through the ANN, but also the shape of the loss landscape, which can directly impact the efficiency with which different types of representations can be learned (Belkin et al., 2019, Du et al., 2019). Second, the success of using the brain’s hierarchical structure to inspire the architecture of ANNs demonstrates that, in principle, it can be beneficial for AI to seek inspiration from the brain when seeking new inductive biases (Hassabis et al., 2017). Importantly, PRC models provide a new way of incorporating biological insights into the design of ANN architectures. If we were to replace the standard units of an ANN with PRC models based on real neurons, this would represent a major change in the architecture of the ANN, one that may provide useful inductive biases.

### Potential advantages of PRC-based inductive biases

When considering the potential inductive biases that PRC models introduce, the natural question is, would these inductive biases actually help or hinder AI? Though it is true that inspiration from brains have provided useful inductive biases for machine learning in the past (LeCun et al., 2015, Hassabis et al., 2017), there is no rule that says that neural phenomena always provide such utility. Indeed, some aspects of physiology may be more related to phylogenetic history and biological constraints than they are to improved learning performance. However, there are a few reasons to think that dendrite inspired PRC models may provide useful inductive biases.

As noted above, research over the last decade has confirmed that network depth is an important architectural consideration for AI (Lee et al., 2017, Li and Sompolinsky, 2020). However, not all depth is equal. Researchers have found that increasing depth is most useful when additional architectural features are included, such as skip connections (Du et al., 2019). If we were to replace the units of an ANN with PRC models based on real neurons that would increase the depth of the network, but in a very particular way. The central question, then, is would the specific form of increased depth that one would obtain from using PRC model units would be helpful?

One reason that PRC models may provide a useful form of depth is that they would help to promote sparsity, which has also been shown to be useful in neural networks (Srivastava et al., 2014). PRC models would help to promote sparsity because they would provide distinct computational subunits with non-linear interactions that could handle different components of a task. For example, it would be possible to have individual dendrites that are responsible for processing distinct types of features, e.g. one set of dendrites for processing facial features, another set for processing body parts, etc. Thus, depending on the input provided, the network could activate only a very sparse set of all the dendrites for processing.

Related to this, there is a growing recognition in the machine learning community that a desirable form of inductive biases for AI would be those that promote the emergence of specialized modules that can be flexibly composed (Goyal et al., 2019, Hinton et al., 2018). This may turn out to be critical to overcoming the limitations of current ANN approaches. Specifically, if an ANN was provided with an architecture that promoted the learning of distinct, specialized modules that can be composed, then it should be possible both to learn more intuitive part-whole relationships that capture the underlying structure of objects in the world more accurately (Goyal et al., 2019, Hinton et al., 2018) and to avoid the catastrophic forgetting that can plague standard ANNs (Masse et al., 2018). However, current systems to promote specialized modules are only loosely inspired by real neural circuits. Therefore, an interesting open question is whether PRC models could provide a good mechanism for implementing brain-inspired inductive biases to promote composability. Notably, dendrites have functionally clustered inputs (Kastellakis et al., 2015), synaptic dynamics can perform LNL operations (Rossbroich et al., 2021), and these phenomena interact (Wilson et al., 2016). Given the fact that PRC models would allow an ANN to learn dendrite-like, flexible, non-linear, recurrent interactions between functionally clustered inputs, it is plausible that they could help with composability. Thus, we would argue that future research should investigate the potential for ANNs that use PRC models inspired by sub-cellular dendritic computation to show better specialization and composability, and less catastrophic forgetting.

## Discussion

In this article, we have illustrated the capabilities of PRC models to capture the various experimentally observed features of dendritic computation, and discussed how this modeling framework may be key to understand the role of dendrites in learning and neuronal computation. In doing so, we have identified modular, composable connectivity motifs (Fig. 2), with the parallel, recurrent and cascade elements forming the basic building blocks of the framework. We have illustrated how the interaction between sodium spikes in the dendrite and the soma (Golding and Spruston, 1998) can be captured using a recurrent motif of LNL subunits (Fig. 4). Furthermore, we have demonstrated that a parallel motif of LNL subunits within a cascade can reproduce the dependence of NMDA spikes on the activation of sodium and calcium channels (Schiller et al., 2000). Finally, we demonstrated that the interaction between back-propagating action potentials and calcium spikes can be captured by a recurrent motif of LNL subunits with a different set of parameters (Fig. 3) (Larkum et al., 1999, 2004). In closing, we have argued that these PRC models of dendritic computation could have an important role in shaping inductive biases (Section h), and thus contribute to the optimization of learning capabilities of the brain.

Our LNL models, however, bear some important limitations. Firstly, the set of operations that are possible within the LNL framework correspond to a subset of the operations that are achievable by the type of dynamical systems used in detailed simulations of dendritic integration (Schiller et al., 2000, Hay et al., 2011, Ujfalussy et al., 2018, Beaulieu-Laroche et al., 2018). For instance, modeling the NMDA spike as two LNL units in parallel followed by another nonlinearity ignores the nonlinear impact that sodium ion channels can have on calcium ion channels via rapid increase of the membrane potential. Another element not captured by the phenomenological LNL model is that ion channel time constants almost always depend on the mean depolarization, which implies an adaptive filter instead of a fixed filter assumed in the LNL model. A second important limitation is that we have assumed that the dendrites remain in a fluctuation-driven regime where the net input is low on average but highly variable. If we were to give strong and sustained inputs to the LNL model, these would saturate the nonlinearity and nonlinear transients would disappear. Such a sustained-depolarization regime has been observed in some experiments (Larkum et al., 2004), but it remains to be seen whether these take place *in vivo* or whether homeostatic mechanisms preserve the fluctuation-driven regime (Vogels et al., 2011). One last limitation of our model is that a very high input variability is capable of making nonlinear operations effectively linear (Ujfalussy et al., 2018). In this case, the complex linear-nonlinear structure operates in a way that can actually be captured by an effective model that is entirely linear. Although some *in vivo* manipulation of dendritic inputs (Xu et al., 2012, Gambino et al., 2014, Doron et al., 2020) argue against this point of view, the full relationship between inputs and outputs of neurons in a naturalistic condition is far from being fully known.

We included in this paper some discussion of the potential machine learning applications of PRC models. As we outlined, there are some reasons to believe that dendritic computations may provide useful inductive biases for machine learning systems. We are hopeful that future research will demonstrate this. However, it also has to be recognized that real dendrites may be solving an implementation problem for neurons, i.e. how can you actually integrate thousands of distinct signals in a physical circuit with space and energy constraints? It is possible that this is the problem which dendrites solve for real neurons, and that dendritic computation is, itself, not important at the algorithmic level. Only by exploring the potential advantages of training AI systems with PRC models inspired by real neurons will we be able to get some initial insight on this mystery.

In summary, our work shows how PRC models can be used to model sub-cellular dendritic computation with a computationally tractable approach. This lays the groundwork for future explorations of the algorithmic implications of dendritic computation, both in the brain, and in machine learning applications. We believe that PRC models will help open the door to exploring the true computational power of dendrites.

## 1. Methods

### 1.1. Linear-nonlinear subunit

The basic component of the modeling framework presented here is the linear-nonlinear subunit, which receives a net time-varying input *I_i_*(*t*) and produces an activation *z*(*t*) as its output

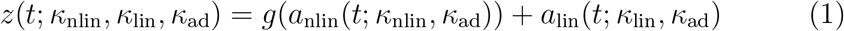

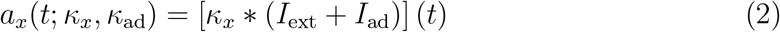

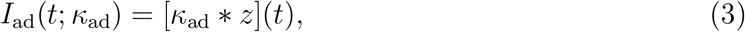

where *g*(·) is a nonlinear activation function; *a*_nlin_(*t*) and *a*_lin_(*t*) are the preactivations for the nonlinear and linear parts of the output, respectively; *I*_ext_(*t*) = ∑_*i*_ *I_i_*(*t*) is the total input from all external sources; *I*_ad_(*t*) is a recurrent adaptation current; and 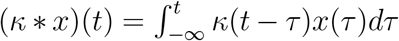 denotes the causal convolution of a filter *κ* with a signal *x* evaluated at time *t*. The subunit is parameterized in terms of the activation filters *κ*_nlin_ and *κ*_lin_, the adaptation filter *κ*_ad_, and the choice of nonlinear activation function *g*(·). Except where noted, all filters in this work are defined as exponential functions

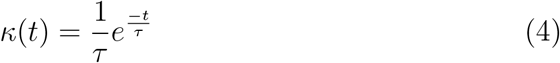

with time constant *τ*. For models with multiple cellular compartments, *I*_ext_(*t*) may include inputs from an external source as well as from other compartments (eg, Methods 1.4). Depending on the model, *z*(*t*), *a*_nlin_(*t*), or *a*_lin_(*t*) may correspond loosely to the voltage of a compartment denoted *V_x_*(*t*) where *x* is the name of the compartment.

### 1.2. Two-compartment model subject to a single dendritic input

The two-compartment model with a recurrent connection from the somatic to the dendritic compartment is defined as follows

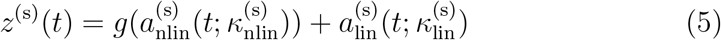

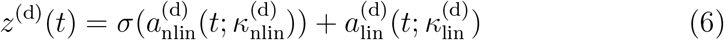

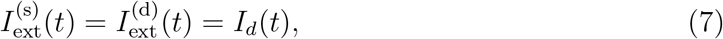

where the dendritic nonlinearity *σ*(·) is the sigmoid function. *g*(·) is the somatic spiking nonlinearity, a function which emits a 1 ms square pulse of amplitude *A* = 2 AU when *z*^(s)^(*t*) crosses the somatic spike threshold from below

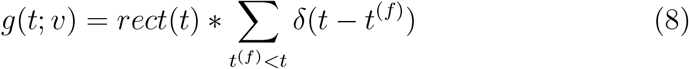

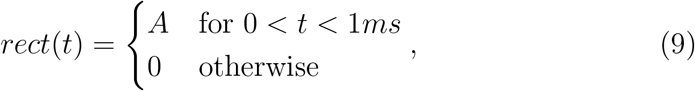

where *t^(f)^* denotes the time of a threshold crossing. Exponential functions with the following time constants were used for the pre-activation filters: 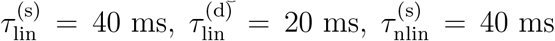, and 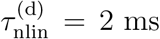. Adaptation filters were set to zero and the terms associated with them were omitted from the above model for simplicity. The activation output *z^(x)^*(*t*) corresponds loosely to the voltage of each compartment; *V_s_*(*t*) ≡ *z^(s)^*(*t*) and *V_d_*(*t*) ≡ *z^(d)^*(*t*) are therefore used to refer to these terms in figures and the main text for ease of interpretation.

### 1.3. Multi-subunit model with parallel processing

The model is composed of three linear-nonlinear subunits which loosely capture the contributions of sodium, calcium, and NMDA voltage-dependent conductances (denoted by the superscripts (1), (2), and (3), respectively) to nonlinear processing of synaptic inputs in a dendritic compartment. Their dynamics are defined as follows

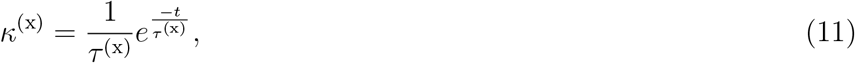

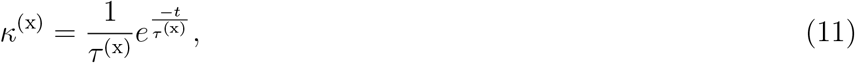

with inputs

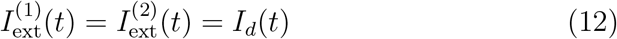

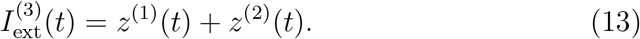

In all three subunits, the non-linear and linear pre-activation filters were set to be equal, such that

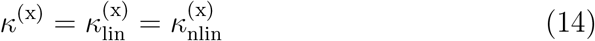

where *τ*^(1)^ = 5 ms, *τ*^(2)^ = 40 ms, and *τ*^(3)^ = 80 ms. The adaptation filters were set to zero and the associated terms dropped for simplicity. The dendritic synaptic-like input current is given by the alpha function

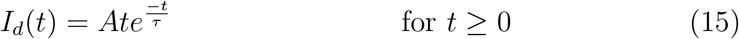

with amplitude *A* and time constant *τ* = 2 ms. The voltage output shown in the figures and main text is analogous to the activation output *V*_x_(*t*) ≡ *z*^(x)^(*t*) for each respective subunit.

### 1.4. Two-compartment model with bi-directional dendro-somatic interactions

The model with bi-directional dendro-somatic interactions is composed of two reciprocally-connected linear-nonlinear subunits (see Section 1.1) as follows

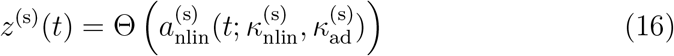

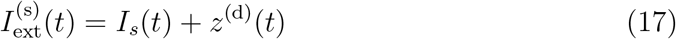

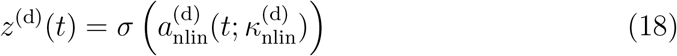

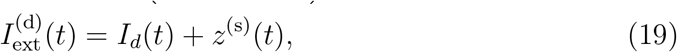

where Θ(·) is the Heaviside step function and *σ*(·) is the sigmoid function. The nonlinear activation filters 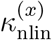 are defined as exponential functions with 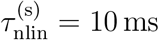 and 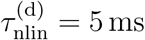. The somatic adaptation filter is defined as

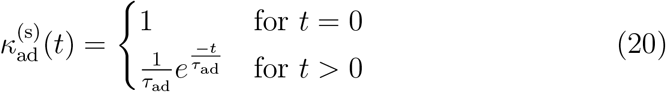

with *τ*_ad_ = 20 ms. In this model, the linear activation filters 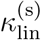 and *κ*_(d)_ are set to zero, along with the dendritic adaptation filter 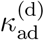. (The terms associated with these filters have been dropped from the above model definition for simplicity.) *I_s_*(*t*) and *I_d_*(*t*) correspond loosely to synaptic inputs to the somatic and dendritic compartments, respectively. The somatic preactivation 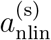 and the dendritic activation *z*^(d)^ loosely correspond to the voltage in their respective compartments. For clarity, we use 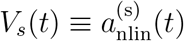 and *V_d_*(*t*) ≡ *z*^(d)^(*t*) to refer to these quantities in the figures and main text.

### 1.5. Multi-layered networks of PRC models

Artificial spiking neural networks with a hidden layer of PRC neurons were constructed following the approach of Refs. (Zenke, 2019, Neftci et al., 2019, Cramer et al., 2020). Briefly, spikes in each layer were integrated as exponentially-decaying synaptic currents in each neuron/PRC-compartment in the following layer, which in turn were integrated by the dynamics of the corresponding PRC subunit as *I*_ext_(*t*). The neurons in the output layer consisted of leaky integrators without threshold or reset. The time-varying voltage of the neurons in the output layer was transformed into a set of class probabilities by applying a softmax operation to the maximum voltage attained by each output unit during each example.

The PRC filters *κ* used in network models were defined as

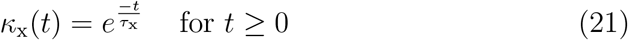

where the membrane time constant *τ*_nlin_ = 10 ms, and the synaptic time constant *τ*_syn_ = 5 ms. Time constants of sodium, calcium, and NMDA filters took the same values as in Methods 1.3. The linear forward filters *κ*_lin_ were dropped for simplicity. The adaptation filter in the somatic compartment was defined as

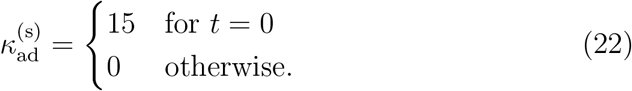

In order to train multi-layered networks of PRC neurons, it was necessary to use a surrogate gradient for the Heaviside step function used to generate spikes in the somatic compartment (since its gradient is zero almost everywhere). Following the approach of (Zenke and Ganguli, 2018, Zenke, 2019), we used the normalized gradient of a sigmoid function to approximate the gradient of the Heaviside function

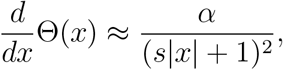

where *α* is a proportionality constant and *s* = 10 is a scale parameter that sets the slope of the sigmoid.

All training was carried out using the Adam optimization algorithm with a learning rate of 0.002 and the negative log likelihood of making a correct class prediction as the loss function.

### 1.6. Numerical methods

Simulations were implemented in Matlab and Python 3.8 using NumPy 1.18.5, SciPy 1.5.0, and ez-ephys 0.4.2. Figures were prepared in Python using Matplotlib 3.2.2, Jupyter 1.0.0, and ez-ephys. Code is available at.

